# *Verticillium dahliae* ubiquitin-specific proteases coordinate key developmental processes and pathogenicity

**DOI:** 10.64898/2026.05.26.727862

**Authors:** Ying-Yu Chen, Miriam Leonard, Melissa Kocatürk, Nils F. Aßmann, Marvin Bromm, Merle Aden, Kerstin Schmitt, Oliver Valerius, Rebekka Harting, Gerhard H. Braus

## Abstract

Ubiquitin is a posttranslational modifier that is conserved among eukaryotes. Ubiquitination alters stability and folding of cellular proteins. Deubiquitinases (DUBs) reverse ubiquitination and often function as part of protein complexes. There are 32 predicted DUB-encoding genes present in the soil-borne phytopathogenic fungus *Verticillium dahliae*. Nuclear ubiquitin-specific protease 3 (Usp3) is a member of the <u>S</u>pt–<u>A</u>da–<u>G</u>cn5 <u>a</u>cetyltransferase (SAGA) complex, whereas Usp1 is predicted to associate with the <u>C</u>OP9 <u>s</u>ig<u>n</u>alosome (CSN), which controls specificities of cellular E3 ubiquitin ligase activities. A proteomics approach using <u>bio</u>tin capture and <u>id</u>entification (BioID) supports that Usp3 regulates gene expression beyond the transcription level. Western experiments showed a dysregulation in ubiquitinated cellular proteins in corresponding deletion strains. Usp3 and Usp1 are both required for fungal development. They regulate microsclerotia formation based on different environmental cues and provide redundant functions in controlling conidiation. Absence of both corresponding genes resulted in significant impairment of conidiospore formation, which is required for fungal propagation within the plant vascular system. This paralysed spreading ability reduced virulence on tomato plants (*Solanum lycopersicum*). In summary, *V. dahliae* responds to environmental cues by Usp3- and Usp1-mediated adjustment of gene expression and protein stability. This is important for key developmental processes of the *V. dahliae* disease cycle and its virulence towards the host plant.

**Author summary:** Ubiquitination and deubiquitination of proteins enable cells to rapidly react to environmental cues and adjust protein stabilities and subsequently transcriptomic profiles. Usp3 is a nuclear deubiquitinase subunit of the <u>S</u>pt–<u>A</u>da–<u>G</u>cn5 <u>a</u>cetyltransferase (SAGA) transcriptional coactivator complex. Usp1 is predicted to be associated with the COP9 signalosome that regulates substrate specificities of the ubiquitin-proteasome system. BioID experiments suggest that other SAGA complex subunits, histone proteins, spliceosomal proteins, ribosomal proteins, a protein that tackles transcriptionally stalled RNAPII, and a protein that degrades mRNA with premature stop codons locate proximal to Usp3 within the cell. Deletion of *USP3* led to the dysregulation of protein ubiquitination. A single deletion of *USP1* did not significantly change ubiquitination profiles, however, a double deletion of *USP1/3* significantly affected the ubiquitin-proteasome system. The altered ubiquitination profile correlated with a dysregulation of key developmental processes. Microsclerotia formation was decoupled from environmental cues in the Δ*USP3* strain, whereas an additional deletion of *USP1* reconnected it in a media-dependent manner. *USP3* and *USP1* contribute to a common governing process in conidiation, and the defect in spreading of the Δ*USP1/3* strain is reflected by a significant reduction in plant pathogenicity.

## Introduction

Posttranslational modifications of proteins are achieved by covalently adding chemical moieties to the amino acid side chains catalysed by specific enzymes, and many of these processes can be enzymatically reversed. Modifications can be phosphorylation, methylation, glycosylation, or acetylation, whereas modifications by proteins of the ubiquitin family lead to ubiquitination, neddylation, or sumoylation of target proteins. Altered protein structure, activity, localisation, interactions, and/or stability of existing proteins facilitates a rapid adaptation to environmental or host-specific cues prior to the adjustment of gene expression profiles (1, 2). Ubiquitin is a posttranslational modifier of 76 amino acids (aa) with a molecular weight of 8.6 kDa. Its C-terminal glycine residue is mainly attached to lysine residues of target proteins through a cascade of E1 ubiquitin-activating enzymes, E2 ubiquitin-transferring enzymes, and E3 ubiquitin ligases (3). All seven lysine residues and the N-terminal methionine of ubiquitin can be additionally modified by further ubiquitin, ubiquitin-like moieties, or by other small molecule modifiers. A target protein can be monoubiquitinated at only one site, multi-ubiquitinated at several sites, or polyubiquitinated by ubiquitin chains of various chain lengths and linkage types (1). Different ubiquitin modifications provide distinct protein surfaces, which can be recognised by effectors of downstream processes. Most ubiquitinated proteins are directed to proteasome for degradation, but ubiquitination can also lead to signalling, or redirect the protein to other cellular sites or compartments (1). Deubiquitinases (DUBs) are enzymes that reverse ubiquitination by hydrolysis. Released ubiquitin then replenishes the ubiquitin pool (4). There are a variety of DUB enzymes that process complex ubiquitin chains and they are classified into six families based on the catalytic mechanism and enzyme domain architectures: <u>u</u>biquitin-<u>s</u>pecific <u>p</u>roteases (USP), <u>u</u>biquitin <u>C</u>-terminal <u>h</u>ydrolases (UCH), <u>o</u>varian <u>tu</u>mor proteases (OTU), <u>M</u>achado–<u>J</u>osephin <u>d</u>omain proteases (MJDs), <u>m</u>otif interacting with ubiquitin-containing <u>n</u>ovel <u>D</u>UB famil<u>y</u> (MINDY), and <u>Ja</u>b1/<u>M</u>PN/<u>M</u>OV34-metalloenzyme (JAMM) (1, 4).

DUBs are often associated with other proteins and form protein complexes that collectively act on target proteins (5). One of the better studied examples is Ubp8 of *Saccharomyces cerevisiae* in the <u>S</u>pt–<u>A</u>da–<u>G</u>cn5 <u>a</u>cetyltransferase (SAGA) complex. SAGA is a transcriptional coactivator complex that is conserved amongst eukaryotes. The *S. cerevisiae* SAGA complex contains a transcription factor-binding module that interacts with transcriptional activators (6), a core module that acts as a structural scaffold to recruit TATA box-binding proteins (7, 8), a histone acetyltransferase (HAT) module that mainly acetylates histone H3 (9), and a DUB module that cleaves ubiquitin from histone H2B (7, 10). Posttranslational histone modifications control the accessibility of specific genomic regions within chromosomes, thereby increase or decrease the expression of certain gene loci epigenetically. The role of the SAGA complex has been studied in several filamentous fungi. The *A. nidulans* SAGA complex coordinates fungal secondary metabolism by acetylating histone H3 of several secondary metabolite gene clusters upon interaction with soil bacteria (11). In the phytopathogenic fungus *Magnaporthe oryzae*, the absence of Ubp8 results in an impaired development, attenuated pathogenicity, and reduced resistance towards oxidative stress (12). Deletion of the gene for the core module Ada1 in *V. dahliae* led to reduced pathogenicity and defects in conidiation and microsclerotia formation (13).

Control of ubiquitination and deubiquitination is provided by the ubiquitin-proteasome system amongst others. The *A. nidulans* COP9 signalosome (CSN) and the cullin-RING ligases (CRLs) are protein complexes that govern ubiquitin-mediated protein degradation (14). CRLs form the largest group of E3 ligases in eukaryotes, and cullin can be posttranslationally modified by the ubiquitin-like protein Nedd8, which leads to an increased degradation of proteins by the proteasome (15, 16). The CSN complex controls ubiquitin homeostasis and affects developmental processes by removing Nedd8 from CRLs and the CSN-associated UspA removes ubiquitin from target proteins (14, 17, 18). The deletion of *uspA* in *A. nidulans* resulted in an increased cellular protein ubiquitination, evoking a disruption of secondary metabolism and organismic development (18).

*Verticillium dahliae* is a haploid soil-borne phytopathogenic fungus with a wide range of host plants of economically importance (19). *V. dahliae* senses and adapts to the different environments during different stages of the infection cycle to ensure successful colonisation (20). *V. dahliae* can remain dormant in the soil for years in the form of heavily melanised microsclerotia (21). Microsclerotia germinate upon encountering environmental cues such as suitable moisture in combination with chemical signals from soil bacteria and host plants (22, 23). The fungal hyphae then grow towards the plant roots and penetrate them to enter the vascular system (19, 24, 25). The fungus then produces asexual conidiospores that spread through the plant and cause plant-wide systemic infection. Typical Verticillium infection symptoms such as wilting, stunting, and vascular discolorations result from a blockage of the vascular transportation system (19). *V. dahliae* then switches from biotrophic to necrotrophic growth in the subsiding phase of the growth season, and forms microsclerotia within the senescing host plant. Microsclerotia are released into the environment along with the decaying plant material (19). Effective agricultural control of Verticillium wilt on economic plants remains challenging due to the resistant microsclerotia in the soil and the limited methods to combat the pathogen once it has entered the host plant (26).

In this study, we aimed to characterise the impact of the *V. dahliae* ubiquitin-specific proteases encoded by *USP3* and *USP1* on coordinating key developmental processes and disease symptom induction, and to establish the *in vivo* proximity labelling with <u>bio</u>tin capture and <u>id</u>entification (BioID) in the plant pathogen *V. dahliae*. *USP3* and *USP1* are homologs of *S. cerevisiae UBP8* and *A. nidulans uspA*, respectively. Our studies revealed that nuclear Usp3 is a member of the *V. dahliae* SAGA complex, and it interacts with ribosomal proteins. Usp3 impacts microsclerotia formation according to the environmental cues, whereas Usp1 plays a minor but overlapping role. Both Usp3 and Usp1 govern conidiation and therefore contribute to disease symptom development in the host plant *Solanum lycopersicum*.

## Results

### The *Verticillium dahliae* JR2 genome encompasses 32 predicted deubiquitinase-encoding genes

BLAST search and searches on corresponding protein domains identified 32 different deubiquitinase (DUB)-encoding genes from five protein families in the *V. dahliae* JR2 genome (S1 Table). These include 15 deubiquitinases from the USP family, five from the UCH family, three from the OTU family, eight from the JAMM family, and one MINDY deubiquitinase. *USP3* and *USP1* are genes predicted to encode ubiquitin-specific proteases in the USP family. *USP3* is orthologous to *Saccharomyces cerevisiae UBP8*, which encodes a member of the deubiquitinase (DUB) that serves as a subunit within the conserved eukaryotic SAGA complex. *USP1* is an ortholog of *Aspergillus nidulans uspA*, whose gene product is associated with the COP9 signalosome. This complex regulates substrate-receptor exchanges for E3 cullin-RING ligases to mediate ubiquitin-labelling of target proteins for 26S proteasome-mediated degradation (27).

*USP3* encodes an ORF of 2097 base pairs (bp), which consists of six exons and seven introns in the coding region (Fig 1a). The encoded protein is 515 amino acids (aa) in length and according to the Ensembl fungi database, it contains a ubiquitin carboxyl-terminal hydrolase domain (PF00443) as well as a Zn-finger in the ubiquitin-hydrolases domain (PF02148) (28). A monopartite nuclear localisation signal (NLS) was predicted by the cNLS mapper server to reside between amino acid positions 144 and 155 of Usp3 (29). We sequenced the coding region of the *USP1* cDNA because of the inconsistency between gene annotations in the databases of the two *V. dahliae* isolates JR2 and Ls.17 (Fig S1). The *USP1* transcript is 7265 bp in length, and the coding region consists of five exons and four introns (Fig 1b). The deduced protein of 1451 amino acids contains a ubiquitin carboxyl-terminal hydrolase domain (PF00443) and a DUSP (domain present in ubiquitin-specific protease) domain (PF6337) (28). Usp1 is predicted to contain a monopartite NLS between amino acid positions 978 and 988 by the cNLS mapper website (29). *V. dahliae* Usp3 and Usp1 share at least 30% sequence similarity with their orthologs in *A. nidulans*, *S. cerevisiae*, and *Homo sapiens* (Fig S2 & S3, S1 Table).

**Fig. 1.**
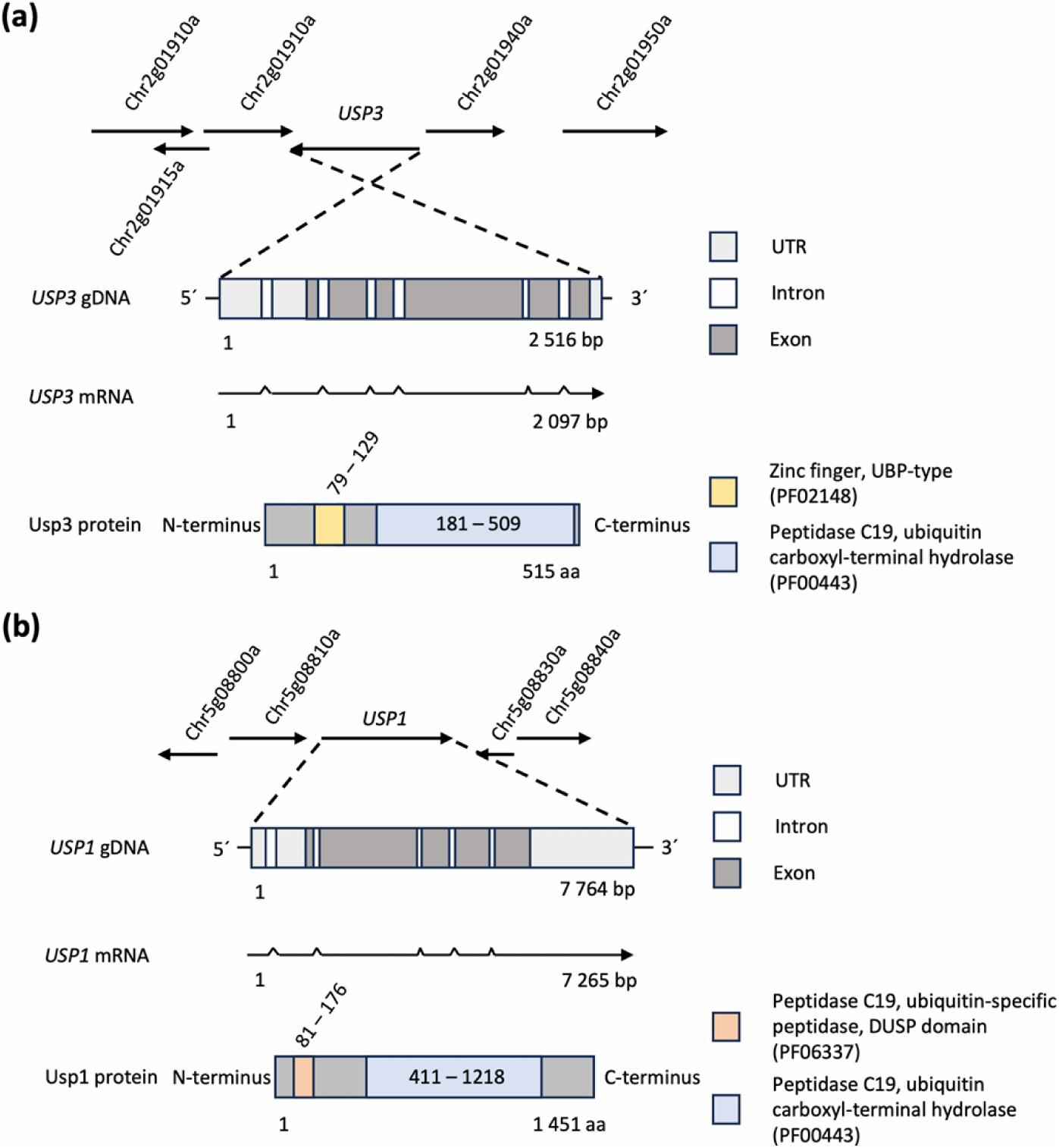
Genomic structure of the *V. dahliae* ubiquitin-specific protease-encoding genes *USP3* and *USP1*. (a) The *USP3* gene contains 6 exons (dark grey) in its coding region, and predicted 5’- and 3’ UTRs (light grey). The gene is transcribed into an mRNA of 2097 bases. The 515 aa Usp3 protein contains a UBP-type zinc finger domain (PF02148; 79 – 129 aa; yellow) and a ubiquitin carboxyl-terminal hydrolase (PF00443; 181 – 509 aa; light blue). (b) The *USP1* gene contains 5 exons in its coding region (dark grey), and predicted 5’- and 3’ UTRs (light grey). The gene is transcribed into an mRNA of 7265 bases. The 1451 aa Usp1 protein contains a ubiquitin-specific peptidase domain (PF06337; 81 – 176 aa; orange) and a ubiquitin carboxyl-terminal hydrolase (PF00443; 411 – 1218 aa; light blue).

### The SAGA complex subunit Usp3 colocalises with nuclear proteins of the chromatin remodelling machinery

As the Usp3 protein was predicted to contain a nuclear localisation signal (NLS), we investigated if the predicted sequence is functional. The in-locus complementation strain expressing GFP-fused Usp3 (*GFP-USP3*) and a strain expressing a GFP-fused biotin ligase that contains the predicted NLS of Usp3 (under the control of the native *USP3* promoter; *TurboID-NLS-GFP*) were transformed to ectopically express RFP-fused histone H2B and observed under the fluorescent microscope. The WT strain expressing RFP-fused H2B and a strain expressing free GFP and RFP-fused H2B served as negative and positive controls. Our results revealed that both TurboID-NLS-GFP and GFP-Usp3 fusion proteins localise to the nucleus, which aligns with the predicted presence of Usp3 in the SAGA transcriptional coactivator complex (Fig 2a).

**Fig. 2.**
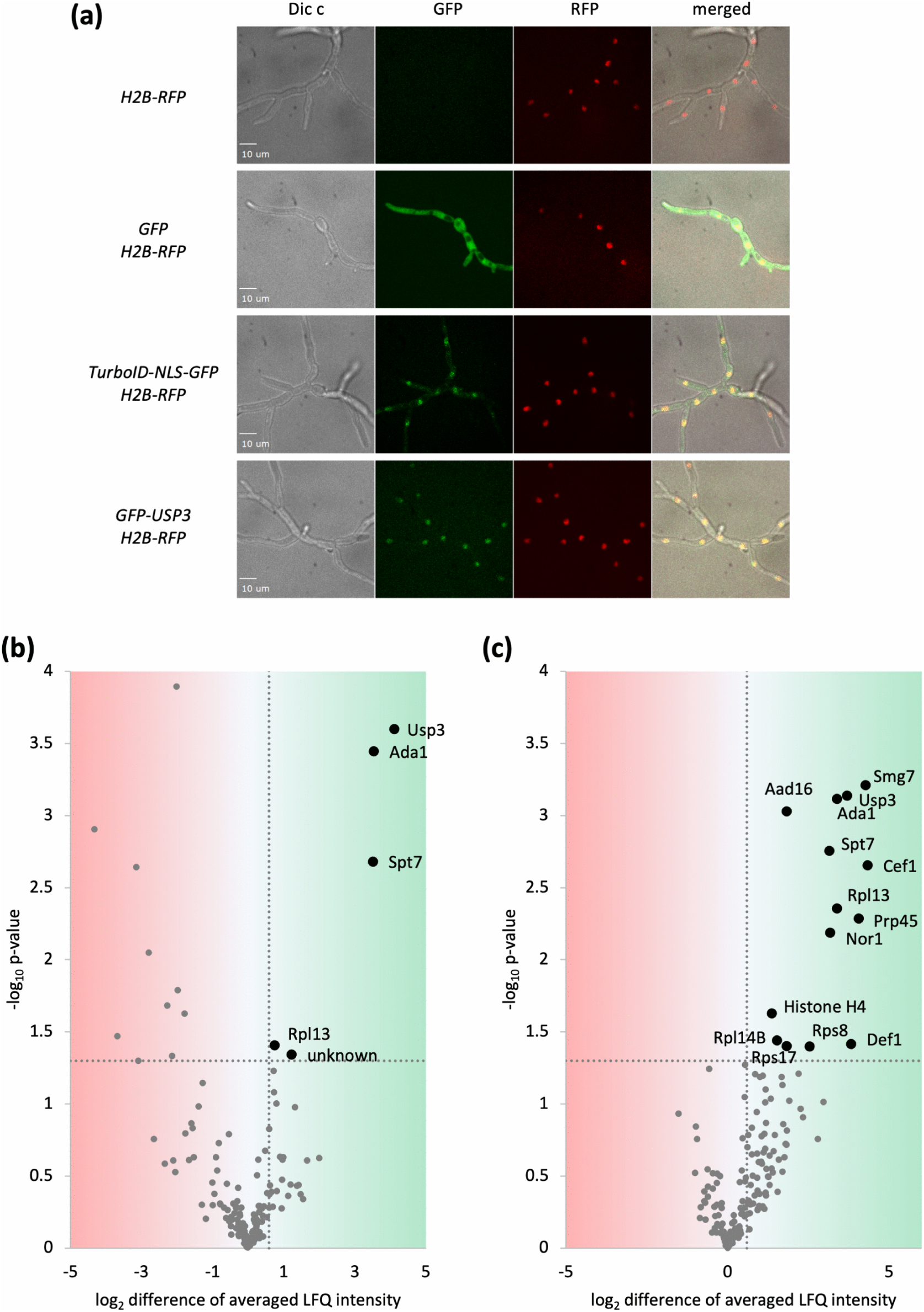
*V. dahliae* Usp3 localises proximal to SAGA complex subunits. (a) Florescence microscopy images were taken from the WT strain, the free GFP strain (*GFP*), the strain expressing GFP-fused to the biotin ligase that contains the NLS of Usp3 under the control of the *USP3* promoter (*TurboID-NLS-GFP*), and the strain expressing GFP-fused Usp3 (*GFP-USP3*). All strains ectopically express RFP-fused histone H2B (*H2B-RFP*). The strains were cultured in liquid PDM overnight, and the images were taken under differential interference contrast view (Dic), green fluorescent protein filter view (GFP), red fluorescent protein filter view (RFP), and a merged view. The white bar at the bottom left corner represents 10 µm. (b/c) Volcano plots of proteins captured and enriched in Usp3-BioID experiments compared to (b) TurboID-NLS-GFP or (c) WT as negative controls. The x-axis depicts the log_2_ transformed differences of averaged LFQ intensities between the experiment groups as indicated. Candidates on the right side of the threshold line are at least 1.5 times more abundant in the TurboID-Usp3 sample (log_2_ difference = 0.585). The y-axis depicts the -log_10_ transformed p-value of the student t-test results, candidates on top of the threshold line (p-value = 0.05) are significantly enriched in either the TurboID-Usp3 or the control samples. The significantly enriched candidates are marked in bold and labelled with their names.

Proteomics BioID experiments were performed to verify the presence of Usp3 in the *V. dahliae* SAGA transcriptional coactivator complex. The expression and biotin ligase activity of TurboID were verified by Western experiments (Fig S4a). All the tested strains expressing the TurboID biotin ligase grew WT-like (Fig S4b). The *TurboID-NLS-GFP* strain and the WT strain served as negative controls for the BioID experiments in order to substract Usp3-independently and non-specifically captured proteins, respectively. Subsequent LC-MS analysis of the captured proteins revealed that the SAGA complex core module members Ada1 and Spt7 were enriched by TurboID-Usp3 against both the *TurboID-NLS-GFP* and the WT control strain (Fig 2b, 2c, Table 1). The 60S ribosomal subunit L13 was also less efficiently captured. Against the less stringent WT control, histone H4, RNA polymerase II degradation factor 1 (Def1), spliceosomal proteins Cef1 and Prp45, nonsense-mediated mRNA decay factor Smg7, additional ribosomal proteins (Rpl14B, Rps17, and Rps8), the putative voltage-gated potassium channel subunit beta-2 (Aad16), and a putative aflatoxin biosynthesis ketoreductase Nor1 were enriched. Altogether, these results suggest that Usp3 is a canonical member of the SAGA complex in *V. dahliae*.

**Table 1.** Significantly enriched proteins in the *V. dahliae* TurboID-Usp3 BioID samples. The predicted functions and localisations of proteins were listed according to the information obtained from the UniProt database.

| Protein IDs | Name | Predicted function | Predicted localisation | $\log_2$ LFQ difference | $-\log_{10}$ Student's t-test p-value | MS/MS counts |
| --- | --- | --- | --- | --- | --- | --- |
| Enriched in TurboID-Usp3 compared to TurboID-NLS-GFP control |  |  |  |  |  |  |
| VDAG_JR2_Ch2g 01930a-00001 | Usp3 | Ubiquitin carboxyl-terminal hydrolase (SAGA complex DUB module) | nucleus | 4.11 | 3.60 | 37 |
| VDAG_JR2_Ch1g 13440a-00001 | Ada1 | Transcriptional coactivator HFI1/ADA1 (SAGA complex core module) | nucleus | 3.53 | 3.44 | 27 |
| VDAG_JR2_Ch1g 01580a-00001 | Spt7 | Transcriptional activator SPT7 (SAGA complex core module) | nucleus | 3.51 | 2.68 | 28 |
| VDAG_JR2_Ch6g 09470a-00001 | Rpl13 | 60S ribosomal protein L13 | ribosome | 0.75 | 1.40 | 42 |
| VDAG_JR2_Ch5g 03480a-00001 | unknown | Isoamyl alcohol oxidase | unknown | 1.22 | 1.34 | 28 |
| Enriched in TurboID-Usp3 compared to WT control |  |  |  |  |  |  |
| VDAG_JR2_Ch5g 06720a-00001 | Smg7 | Nonsense-mediated mRNA decay factor | nucleus | 4.27 | 3.21 | 137 |
| VDAG_JR2_Ch2g 01930a-00001 | Usp3 | Ubiquitin carboxyl-terminal hydrolase (SAGA complex DUB module) | nucleus | 3.69 | 3.14 | 37 |
| VDAG_JR2_Ch1g 13440a-00001 | Ada1 | Transcriptional coactivator HFI1/ADA1 (SAGA complex core module) | nucleus | 3.38 | 3.11 | 27 |
| VDAG_JR2_Ch1g 12360a-00001 | Aad16 | Voltage-gated potassium channel subunit beta-2 | unknown | 1.82 | 3.03 | 288 |
| VDAG_JR2_Ch1g 01580a-00001 | Spt7 | Transcriptional activator SPT7 (SAGA complex core module) | nucleus | 3.14 | 2.75 | 28 |
| VDAG_JR2_Ch8g 03990a-00001 | Cef1 | Pre-mRNA-splicing factor cef-1 | cytosol, nucleolar peripheral inclusion body, post-mRNA release spliceosomal complex, Prp19 complex | 4.33 | 2.65 | 152 |
| VDAG_JR2_Ch6g 09470a-00001 | Rpl13 | 60S ribosomal protein L13 | ribosome | 3.39 | 2.35 | 42 |
| VDAG_JR2_Ch1g 12060a-00001 | Prp45 | Pre-mRNA-processing protein 45 | nucleus | 4.06 | 2.28 | 80 |
| VDAG_JR2_Ch8g 10540a-00001 | Nor1 | Aflatoxin biosynthesis ketoreductase nor-1 | cytoplasm | 3.17 | 2.19 | 47 |
| VDAG_JR2_Ch5g 07535-00001; VDAG_JR2_Ch5g 05710a-00001 | Histone H4 | Histone | nucleus | 1.36 | 1.63 | 20 |
| VDAG_JR2_Ch2g 12510a-00001 | Rpl14B | 60S ribosomal protein L14-B | ribosome | 1.52 | 1.44 | 23 |
| VDAG_JR2_Ch8g 11230a-00001 | Def1 | RNA polymerase II degradation factor 1 | nucleus, cytoplasm | 3.82 | 1.41 | 606 |
| VDAG_JR2_Ch8g 04220a-00001 | Rps17 | 40S ribosomal protein S17 | ribosome | 1.83 | 1.40 | 31 |
| VDAG_JR2_Ch4g 02540a-00001 | Rps8 | 40S ribosomal protein S8 | ribosome | 2.54 | 1.40 | 60 |

### Usp3 contributes to the homeostasis of cellular protein ubiquitination

Western experiments were performed using a primary antibody that recognises ubiquitin to investigate the impact of Usp3 and Usp1 in regulating cellular protein ubiquitination. The absence of *USP1* did not result in a statistically significantly difference in ubiquitinated proteins compared to the WT signal. However, deletion of *USP3* resulted in a significant reduction of ubiquitinated proteins by 38%, whereas proteins from the Δ*USP1/3* double deletion strain showed 54% higher rates of ubiquitination (Fig 3). These results suggest that Usp1 may share overlapping functions with other deubiquitinases in *V. dahliae* as the single deletion did not result in a statistically significant change in the amount of ubiquitinated proteins. The absence of *USP3* resulted in a dysregulation of ubiquitinated proteins, and the removal of *USP1* in the Δ*USP3* background resulted in a much greater increase in ubiquitinated proteins.

**Fig. 3.**
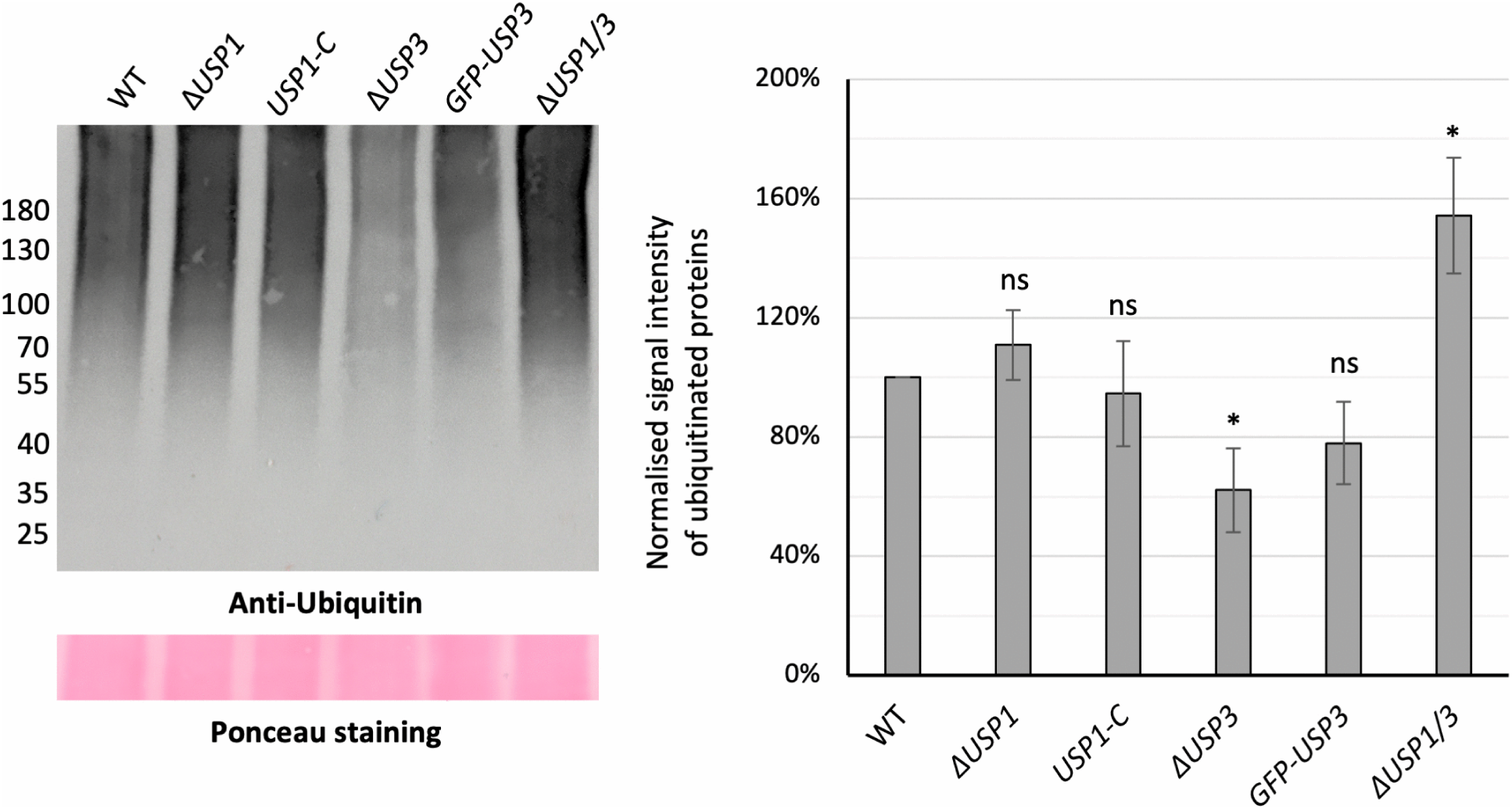
The ubiquitination profile of *V. dahliae* cellular proteins is affected by both Usp3 and Usp1. The single deletion of *USP3* resulted in a decreased amount of ubiquitinated cellular proteins, whereas the single deletion of *USP1* did not result in a statistically significant difference in ubiquitinated proteins. The double deletion of *USP3* and *USP1* resulted in a increased amount of ubiquitinated cellular proteins. The WT, the *USP1* and *USP3* single deletion strains (Δ*USP1* and Δ*USP3*), the *USP1* and *USP3* complementation strain (*USP1-C* and *GFP-USP3*), and the *USP1* and *USP3* double deletion strain (Δ*USP1*/*3*) were cultured in liquid PDM for 6 days before the mycelia were harvested for protein extraction. Western experiments were performed with antibody recognising ubiquitin (Anti-Ubiquitin). The positions of the protein marker are marked on the left, and Ponceau staining served as loading control to ensure that an equal amount of protein was loaded in each lane. Signal intensity of each lane was quantified and normalised by the signal intensity of the WT strain in the same biological replicate. One sample t-test was performed for statistical analysis (*, p < 0.05; ns, not significantly different).

### The Usp3 deubiquitinase links *V. dahliae* microsclerotia development to environmental cues

The role of Usp3 and Usp1 during *ex planta* growth was examined by point inoculating equal amount of conidiospores on minimal medium (CDM) plates and media that favour microsclerotia formation such as pectin-rich simulated xylem medium (SXM) and minimal medium with cellulose as carbon source (CDM + cellulose). An increased microsclerotia formation was observed on CDM plates in the absence of Usp3, whereas the *USP3* single deletion strain was less melanised on media that typically induce resting structure formation. Microsclerotia formation was not substantially different from the WT strain when only *USP1* was deleted. The Δ*USP1/3* double deletion strain had a less severe melanisation phenotype on SXM and CDM, but further reduced melanisation when cultured on CDM with cellulose (Fig 4). These results suggest that both *USP3* and *USP1* are involved in regulating microsclerotia formation, but the effect of *USP3* is more dominant. The absence of Usp3 uncouples microsclerotia development from environmental cues. Phenotypes observed in the double deletion strain imply that Usp1 and Usp3 react to different nutritional cues, but they have overlapping functions in microsclerotia formation under certain conditions.

**Fig. 4.**
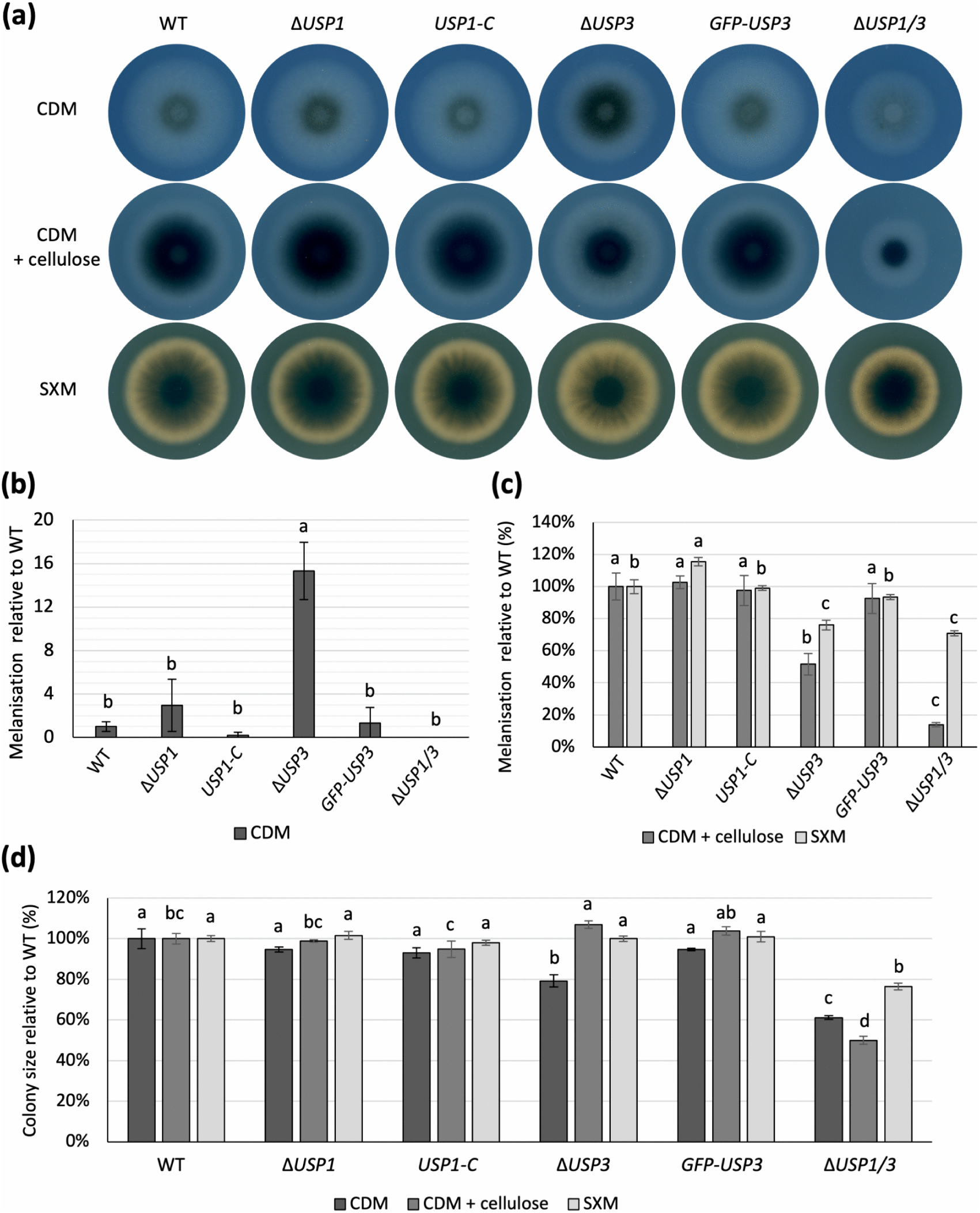
Usp3 and Usp1 regulate *V. dahliae* microsclerotia formation. (a) Less mircrosclerotia are formed on plates that favour resting structure formation (SXM and CDM + cellulose) when *USP3* is deleted, whereas more mircrosclerotia are formed on CDM plates when *USP3* is deleted. The Δ*USP1*/*3* strain further reduced microsclerotia formation on CDM + cellulose plates, but partially restored WT-like melanisation phenotype on SXM and CDM plates. All tested strains were point inoculated on 30 mL agar plates and incubated at 25°C for 10 days. The bottom view of each plate is shown. Melanisation levels of colonies spotted on CDM quantified and presented in (b), and melanisation levels of colonies grown on pectin-rich-SXM and CDM with cellulose as carbon source are quantified and presented in (c). Melanisation levels are normalised by the average of the WT in the respective culturing medium. (d) The size of each colony is quantified and normalised by the average size of the WT colony size in the respective medium. Statistical analyses were performed by one-way ANOVA with post-hoc Tukey HSD test. A difference of the lower-case letter on top of each bar indicates significant difference (p < 0.05).

Secondary metabolites were extracted from CDM or SXM plates cultured with WT or the Δ*USP3* strain. Empty plates served as media control. The Δ*USP3* strain showed no observable differences in its metabolite profile compared to the WT control although a reduction in melanisation was observed in the *USP3* mutant strains in the *ex planta* phenotypic analysis (Fig S5).

### Usp3 and Usp1 deubiquitinases have redundant functions in fungal conidiation and pathogenicity

The resistance of microsclerotia and the rapid spread of conidiospores in the plant vascular system are two major factors that make Verticillium wilt difficult to control (19). To examine whether *USP3* or *USP1* are involved in the regulation of conidiation, equal numbers of conidiospores of the tested strains were inoculated in 50 mL SXM and incubated at 25°C. The produced conidiospores were harvested and counted five days post-inoculation. Conidiospores produced by the *USP3* and *USP1* single deletion strains were not significantly different from the WT strain, but the counts of conidiospores significantly decreased in the *USP3* and *USP1* double deletion strain (Fig 5a). *In planta* phenotypes of the *USP3* and *USP1* deletion strains were studied by inoculating ten-day-old tomato seedlings with equal numbers of conidiospores. Disease scores of the infected plants were evaluated three weeks post-inoculation according to the relative fresh weight, highest vegetative point, longest leaf, and vascular discolouration compared to the MOCK-treated plants. An increased number of healthy plants was recorded from the ones inoculated with the *USP1* single deletion strains compared to WT, and disease symptoms were further reduced when the plants were inoculated with the *USP3* and *USP1* double deletion strain (Fig 5b). Our observations suggest that *USP3* and *USP1* may be redundant in the regulation of conidiation. The paralysed ability of the *USP3* and *USP1* deletion strains to spread through conidiospores is reflected on the compromised severity of disease symptoms.

**Fig. 5.**
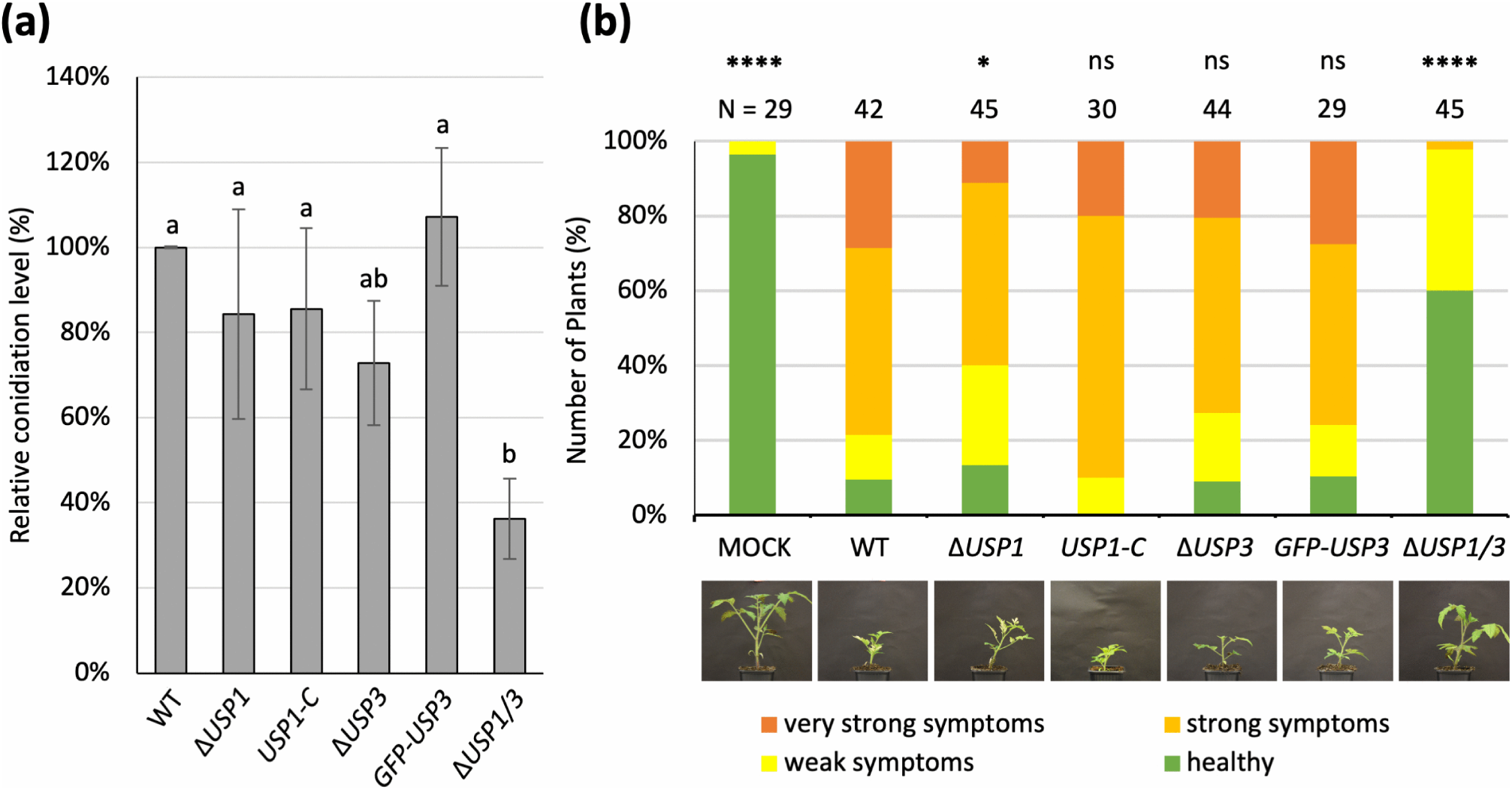
Usp1 and Usp3 affect *V. dahliae* pathogenicity and conidiation. (a) Usp1 and Usp3 deubiquitinases have redundant functions in regulating conidiation. The conidiation levels of the Δ*USP1*/*3* strain was less than 40% of the WT level. Three biological replicates were performed, and statistical analysis to compare the relative conidiation level of each strain was done by one-way ANOVA with post-hoc Tukey HSD test. A difference of the lower-case letter on top of each bar indicates significant difference (p < 0.05). (b) *USP1* and *USP3* have redundant functions in the regulation of virulence. Plants treated with the *USP1* and *USP3* double deletion strain are healthier than WT treated plants, with heavier fresh weight, taller highest vegetative point, longer leaves, and less vascular discolouration. Tomato seedlings were inoculated with water (MOCK), the WT, the *USP1* and *USP3* single deletion strains (Δ*USP1* and Δ*USP3*), the *USP1* and *USP3* complementation strains (*USP1-C* and *GFP-USP3*), and the *USP1* and *USP3* double deletion strain (Δ*USP1*/*3*). Representative photos of tomato plants treated by each strain are shown below. The disease scores were assigned to each plant three weeks post-inoculation according to the relative growth to the MOCK-treated plants. The number of evaluated plants is listed above, and statistical analysis were performed by Mann-Whitney U test (p < 0.0001, ****; p < 0.05, *; not significantly different, ns).

## Discussion

Posttranslational modifications allow the flexibility for the cell to quickly react to the changing environment by varying interaction surfaces. This results in the adaptation of folding, localisation, or stability of distinct cellular proteins. The diverse possibilities of ubiquitination patterns enable the cell to interpret the ubiquitin code and react accordingly. Ubiquitination can be reversed by deubiquitinating enzymes (DUBs). Many DUBs exist in the form of protein complexes to act on common targets. Examples include the <u>S</u>pt–<u>A</u>da–<u>G</u>cn5 <u>a</u>cetyltransferase (SAGA) complex member Ubp8 in *S. cerevisiae*, and the COP9 signalosome (CSN) associated UspA in *A. nidulans* (10, 18). How ubiquitination and deubiquitination affect the growth and pathogenicity of the phytopathogenic fungus *Verticillium dahliae* remains understudied. This study focuses on *USP3* and *USP1*, which are *V. dahliae* homologs of *S. cerevisiae UBP8* and the *A. nidulans uspA* respectively. In *S. cerevisiae*, the substitution of the histone H2B ubiquitination site or the deletion of *UBP8* both resulted in lowered transcript levels of SAGA complex regulated genes (10). Histone H2B deubiquitination happens in yeast or human cells upon UV irradiation, which results in DNA damage and RNA polymerase II (RNAPII) stalling (30). The absence of *ubp8* in yeast cells leads to reduced nucleotide repair and increased RNAPII degradation (30). In *V. dahliae*, histone H2B can be ubiquitinated by the E3 ligase Bre1, which is the only studied ubiquitinase in the fungus (31). VdBre1 affects *V. dahliae* pathogenicity by regulating fungal lipid metabolism, secondary metabolism, and penetration structure formation (31, 32).

In this study, we investigated the cellular localisation and the local environment in the proximity of Usp3. The role of Usp3 in the nucleus was supported by the functional NLS and the identification of SAGA complex core module members Ada1 and Spt by BioID experiments (Fig 2). Although it is known in *S. cerevisiae* that Ubp8 removes ubiquitin from histone H2B, we were only able to identify histone H4 but not histone H2B in the proximity of Usp3. This is likely due to the trypsin digestion method used during LC-MS sample preparation, as it makes the detection of lysine-rich histones difficult during LC-MS measurements. In addition to potentially altering the transcription of genes through SAGA-complex-mediated histone modifications, our BioID results suggested that the *V. dahliae* Usp3 and potentially the SAGA complex is more widely involved in other processes of gene expression regulation. Usp3 was also found to be in proximity of the RNAPII degradation factor 1 (Def1), which is known to mediate the removal of stalled RNAPII on damaged DNA sequences (33, 34). A transcribed pre-mRNA is typically processed by the spliceosome to remove introns (35), and faulty mRNAs that contains a premature stop codon will be detected and degraded by the nonsense-mediated mRNA decay (NMD) pathway to avoid truncated or non-functional proteins accumulating in the cell (36). The yeast homologs of the Usp3 interaction partners Cef1 and Prp45 are associated with the spliceosome (37), whereas the interaction partner Smg7 is involved in the NMD pathway (38). Notably, several ribosomal proteins were also discovered in the Usp3 microenvironment. Ribosomal proteins typically enter the nucleus during ribosome biogenesis, during which they concentrate in the nucleolus to assemble immature ribosomal subunits (39). Having ribosomal proteins nearby Usp3 suggests that Usp3 might also be regulating gene expression at the translation level by controlling the availability of ribosomes. In summary, the presence of key proteins that remove stalled RNAPII, spliceosome members, and an NMD pathway member implies that the *V. dahliae* SAGA complex might play a role in a coordinated gene expression control mechanism that mitigates stress conditions.

Our results revealed that both Usp3 and Usp1 control the amount of ubiquitinated proteins within the cell. However, the deletion of *USP3* resulted in a decreased instead of increased amount of ubiquitinated proteins. To date, histone H2B is the best-known target of *S. cerevisiae* Ubp8 (10), and our BioID results also support that *V. dahliae* Usp3 is mostly involved in gene expression regulation at the transcriptional level (Tabel 1). The reduction of ubiquitinated proteins may be due to the reduced transcription of E1 ubiquitin-activating enzymes, E2 ubiquitin-transferring enzymes, or E3 ubiquitin ligases, which results in a paralysed ubiquitin-proteasome system (UPS) and fewer ubiquitinated proteins. In contrast to Usp3/Ubp8, the *A. nidulans* UspA is known to be associated with the COP9 signalosome that regulates the UPS at the protein level (18). Other *V. dahliae* deubiquitinases may share common targets as Usp1, which results in a slight and statistically insignificant increase in ubiquitinated proteins. However, the removal of *USP1* in a Δ*USP3* background may have resulted in the malfunctioning of the COP9 signalosome-regulated feedback loop in a cell that already lacks a functioning UPS, resulting in an increased amount of ubiquitinated protein.

The inability of the Δ*USP3* and Δ*USP1/3* mutant strains to maintain homeostasis of ubiquitinated proteins is reflected on their ability to form microsclerotia upon receiving certain environmental cues. *V. dahliae* typically forms the heavily melanised microsclerotia when growth conditions are no longer favourable, for example when the hosts growth season ends (19). The pectin-rich SXM and a minimal medium supplemented with cellulose (CDM + cellulose) are commonly used culture conditions in *V. dahliae* research. These media are thought to approximate aspects of plant cell wall-derived polysaccharides, and they support microsclerotia formation under laboratory conditions (40, 41). In the phytopathogen *M. oryzae*, the deletion of *Ubp8* resulted in irregular melanisation (12). Our experiments showed that *V. dahliae* forms less microsclerotia in resting structure-inducing media but has an increased amount of microsclerotia in sucrose-containing minimal medium (CDM) in the absence of *USP3* (Fig 4). The dysregulation of microsclerotia formation indicates that *V. dahliae* has lost its ability to correctly react to nutrient cues upon the deletion of *USP3*. Although the effect of the predicted COP9 signalosome-associated Usp1 on melanisation was minor when Usp3 is present, its role in resting structure formation is evident in the Δ*USP1/3* strain. The deletion of *USP1* in the Δ*USP3* strain rescued the melanisation phenotype in SXM and CDM, but further reduced melanisation in CDM with cellulose as carbon source. The Usp3-coordinated gene expression control mechanisms have a more profound influence on microsclerotia formation than the effect of Usp1-regulated proteasome-mediated protein degradation, and although the two DUBs might have redundant functions under certain nutrient environments, they can also react differently to different nutrient sources. Our results suggest that the microsclerotia formation may be regulated differently when pectin is still readily available from senescing plant material, compared to when the underlying cellulose microfibrils are also exposed for breakdown (42).

A successful Verticillium wilt disease development depends on the coordination of developmental processes, mitigation to the stress conditions caused by the host defence mechanisms, and counteracting hostile environments by secreted proteins or metabolites (20, 43). The deletion of *USP1* led to reduced pathogenicity, and even more healthy plants were observed when both *USP1* and *USP3* are deleted (Fig 5b). The less severe disease symptoms can be mainly explained by the paralysed ability to undergo conidiation, which is essential for this vascular pathogen to rapidly spread in the plant (Fig 5a). As a significant defect in conidiation was only observed in the Δ*USP1/3* strain but not from the single deletion strains, it is possible that the *USP3* and the SAGA complex-controlled conidiation genes encode conidiation-related proteins that are also targeted by Usp1. Although it is known in *A. nidulans* that the SAGA complex HAT module regulates secondary metabolism (11), we were unable to detect any differences between metabolites extracted from the Δ*USP3* strain and the WT strain in our tested conditions (Fig S5). Unlike the effect of acetylation in the chromatin-based regulation of fungal secondary metabolism, the role of the SAGA complex DUB module in fungal secondary metabolism remains unclear.

In summary, our findings support a wider role of *V. dahliae* Usp3 in gene expression regulation than just the SAGA complex-mediated transcriptional control. Usp3 is presumably part of the coordination of other stress-induced transcriptional regulatory processes, mRNA processing as well as ribosome biogenesis. We also elucidated the role of deubiquitinases in the regulation of key developmental processes such as conidiation and resting structure formation (Fig 6). Since the rapid spread of conidiospores in the vascular system and the resistance of the microsclerotia remains as the main obstacles that prevent effective disease control, this study presents two promising potential targets for future efforts to combat Verticillium wilt.

**Fig. 6.**
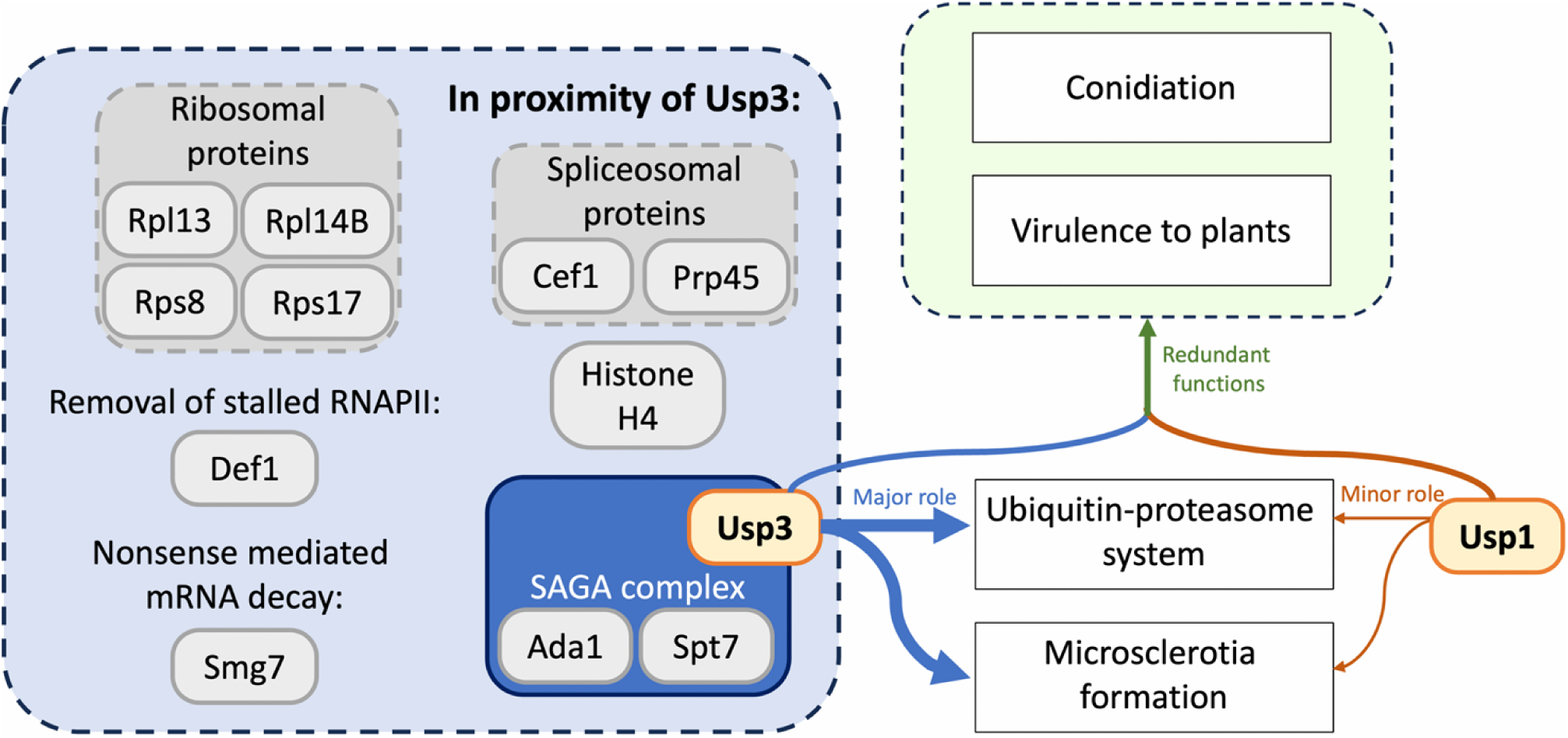
Proteins in the proximity of *V. dahliae* deubiquitinase Usp3 and the functions of Usp3 and Usp1 in the plant pathogen. Fungal Usp3 is a member of the SAGA transcriptional regulator complex (blue box, solid outline), whereas Usp1 is anticipated to be involved in the ubiquitin-proteasome system. Proximity labelling experiments by BioID revealed that Usp3 may be involved in a wider range of functions than the currently known SAGA complex function as a transcriptional regulator. In addition to Histone and SAGA complex subunits, spliceosomal proteins Cef1 and Prp45, RNAPII degradation factor Def1, and Smg7 involved in nonsense-mediated mRNA decay were significantly enriched by Usp3 (blue box, dashed outline). Usp3 and Usp1 shares a redundant function in regulating conidiation and virulence towards plants (green arrow). Both Usp3 and Usp1 are involved in the ubiquitin-proteasome system and the regulation of microsclerotia formation, but the effect of Usp3 (thick blue arrow) is more dominant than Usp1 (thin orange arrow).

## Materials and methods

### Strains and growth conditions

*Verticillium dahliae*, *Escherichia coli*, and *Agrobacterium tumefaciens* strains used in this study are listed in S2 Table. Bacterial cultures were grown in lysogeny broth (LB) medium supplemented with 100 µg/mL kanamycin (AppliChem) when necessary (44). *V. dahliae* cultures were grown in simulated xylem medium (SXM), potato dextrose medium (PDM), or Czapek Dox medium (CDM) as previously described (45). *E. coli* and *A. tumefaciens* samples were incubated at 37°C and 25°C respectively, whereas the *V. dahliae* samples were incubated at 25°C.

### Bioinformatic tools and verification of gene annotation

Gene annotations were obtained from the Ensembl Fungi database (28). Nuclear localisation signals were predicted by the cNLS mapper website with default settings (29). Due to the inconsistencies between *USP1* annotations in the *V. dahliae* JR2 and the *V. dahliae* Ls. 17 databases, the cDNA of *USP1* was sequenced to verify its annotations. The coding region of the *USP1* cDNA was amplified by primer pair RH760 and RH765. The PCR product were ligated into the pJET1.2/blunt cloning vector by the CloneJET PCR Cloning Kit (Thermo Fisher Scientific) and sequenced by Microsynth Seqlab (Göttinegn, Germany). Predicted functions and localisation of proteins enriched in the BioID experiments were obtained from the UniProt database (46).

### *Verticillium* mutant strain construction

Sequence confirmed plasmids that contain the deletion or complementation cassette of *USP1* or *USP3* were transformed into to *A. tumefaciens* AGL1 by the heat shock method (47). *V. dahliae* strains were transformed by *Agrobacterium*-mediated transformation as described in previous studies (24). The genome of the generated *V. dahliae* mutant strains were confirmed by Southern hybridisation (Fig S6). Plasmids and primers used in this study are listed in S3 and S4 Tables.

### Protein extraction and Western experiments

For Western experiments to analyse ubiquitinating patterns, 1 x 10^6^ conidiospores of each tested fungal strain were cultured in 50 mL liquid PDM, and protein samples were extracted from grinded mycelia by modified denaturing lysis buffer as previously described (43). The ubiquitin-antibody (clone P4D1-A11, Merck Millipore) antibody was used as primary antibody in a 1:2000 dilution for Western experiments.

For Western experiments to analyse biotinylating patterns and BioID experiments, conidiospores of each tested fungal strain were inoculated in 50 or 500 mL liquid PDM to reach a final concentration of 1 x 10^6^ conidiospores per mL. The cultures were incubated at 25°C for three days, and 410 nM of biotin was supplemented to the cultures 30 min before mycelial harvest. Protein samples were extracted from grinded mycelia by modified denaturing lysis buffer according to the protocol described in previous studies (43). For Western experiments to analyse biotinylating patterns, the 1:5000 diluted Pierce High Sensitivity Streptavidin-HRP (Thermo Fisher Scientific), 1% BSA, and 0.1% Tween were solved in 1x PBS and incubated with the Western membrane for 1 hr. For Western experiments to analyse the expression of the biotin ligase (TurboID), BirA-antibody (NBP2-59939; Bio-Techne) was diluted 1:2000 in TBS containing 5% milk powder.

All samples were cultured at 25°C with shaking at 125 rpm for the described incubation time. Protein concentration was determined by Bradford assay, and Western experiments were performed as described in earlier studies with modified primary antibodies as stated (45, 48). Loading control for Western experiments was done by Ponceau staining.

### Confocal microscopy

The localisation of GFP, TurboID fused C-terminally with the nuclear localisation signal of Usp3 and GFP (TurboID-NLS-GFP), as well as the GFP-Usp3 fusion protein were visualised by confocal fluorescent microscopy with GFP filters. Histone H2B fused with N-terminus RFP was ectopically expressed and observed by confocal fluorescent microscopy with RFP filters to visualise the nuclei (49).

### *In vivo* proximity labelling with biotin (BioID)

BioID experiments were conducted by expressing the *Sordaria macrospora* codon-optimised TurboID in *V. dahliae* (50). Three biological replicates were performed according an earlier described protocol with minor modifications (50). In brief, equal amount of total proteins were denatured at 65°C for 5 min and the biotinylated proteins were enriched by Strep-Tactin Sepharose (IBA Lifesciences GmbH) as described (50), but the eluted biotinylated protein samples were concentrated by chloroform methanol precipitation (51). Protein samples were resuspended by 1x loading dye (0.25 M Tris-HCL pH6.8, 15% β-mercaptoethanol, 30% glycerol, 7% SDS, 0.3% bromophenol blue) dissolved in TE buffer and loaded onto 12% polyacrylamide gels. The gels were incubated in fixing solution for at least 45 min before being stained by colloidal Coomassie blue for visualisation. The lane that contained the protein samples were fractioned into gel pieces, de-stained by fixing solution, and washed by deionised water. In-gel trypsin digestion, C-18 STAGE tip purification, and LC-MS analysis were performed as described (52). The raw LC-MS data was analysed by MaxQuant 1.6.10.43 and Perseus 1.6.0.7 with the *V. dahliae* JR2 protein database of Ensembl Fungi (28, 53, 54). Parameters of the analysis were set according to earlier studies, but with biotinylation added to the variable modifications during MaxQuant analysis (48, 52).

### Phenotypic analysis

5 x 10^4^ of freshly harvested conidiospores of the tested *V. dahliae* strains were point inoculated on 30 mL SXM agar, CDM agar, and CDM agar plates with sucrose substituted with 3% cellulose as previously described (43). The plates were incubated at 25°C, and the bottom view of each plate was documented 10 days post inoculation (dpi). Melanisation levels were quantified according to the methods described in an earlier study (43).

### Secondary metabolite extraction and analysis

Two CDM or SXM plates were inoculated with 1 x 10^6^ of freshly harvested conidiospores of each tested strain. The plates were incubated at 25°C for two weeks, and the metabolites were extracted from homogenised agar as previously described (48). The metabolites were detected by a Q Exactive Focus Orbitrap mass spectrometer coupled with an UltiMate 3000 high-performance liquid chromatography (HPLC) and a CAD-3000 (Thermo Fisher Scientific) (55). The chromatograms were analysed by FreeStyle 1.6 software (Thermo Fisher Scientific).

### Quantification of conidiation

To quantify the conidiation ability of each tested strains, 2 x 10^5^ of freshly harvested conidiospores were inoculated into 50 mL of liquid SXM. Four technical replicates were performed for each biological replicate, and the cultures were incubated at 25°C for 5 days with shaking at 125 rpm. The conidiospores were quantified and normalised as described by Starke et al. (45). Three biological replicates were performed, and statistical analysis was carried out by one-way ANOVA with post-hoc Tukey HSD test.

### Tomato plant infection

Infection experiments on 10-days old *Solanum lycopersicum* seedlings were performed as described (48), and the disease scores of each plant were evaluated according to the descriptions of Starke et al. (45).

## Acknowledgments

We thank N. Scheiter for technical assistance, Dr. J. Starke, Dr. A. Nagel, Dr. A.M. Köhler, and Dr. A. Strohdiek for the discussions and support.

## Supporting information

**S1 Fig. The *USP1* cDNA contains 5 exons and is 4356 bp in length.**

**S2 Fig. Amino acid sequence of Usp3 and their paralogs.**

**S3 Fig. Amino acid sequence of Usp1 and their paralogs.**

**S4 Fig. Expression of the functional TurboID in *V. dahliae* does not affect growth.**

**S5 Fig. The absence of *USP3* did not change the secondary metabolite profile in the tested conditions.**

**S6 Fig. Verification of the *V. dahliae USP3* and *USP1* mutant strains.**

**S1 Table. List of predicted deubiquitinases in *V. dahliae* and their paralogs in *A. nidulans*, *S. cerevisiae*, and *Homo sapiens*.**

**S2 Table. Bacterial and fungal strains used in this study. S3 Table. Plasmids used in this study.**

**S4 Table. Primers used in this study.**

